# Novel gramicidin-derived *β*-helical antimicrobial peptides

**DOI:** 10.1101/2023.12.31.573784

**Authors:** Edward Lambden, Mark I. Wallace, Martin B. Ulmschneider

## Abstract

Combating the threat of multidrug resistance bacteria requires many different approaches. Using structural motifs and sequences from existing antibacterial compounds affords a fresh attack against such bacteria, while circumventing inherited resistance against the existing compound. Gramicidin is a 16 amino acid peptide with an alternating l- and d-motif with highly bacteriocidal effects in gram-positive membranes. In the membrane it assumes a *β*-helix form with two peptides aligned by their N-terminal interactions, and only conducts in such a form. Using gramicidin as a basis, this study employs molecular dynamics simulations as an initial design step and evaluates gramicidin-derived candidates based on their backbone hydrogen-bond interactions, capacity for conductance, profile of their RMSD over the course of the simulation, and how the peptide interacts with the phospholipid headgroups. There are twelve candidate systems with results comparable to gramicidin and the factors at play in their efficacy are examined.

## 1 INTRODUCTION

*β*-helices have been a subject of scientific interest for much of the last century, as is the case for the linear peptide gramicidin (1), a family whose antibacterial properties have been known and investigated since the 1940s (2) (3). It is expressed in the soil bacteria *Bacillus brevis*, with its name derived from its bactericidal effects against Gram-positive bacteria. Gramicidins have a hydrophobic amino acid sequence of: Formyl-X_1_-GLY-ALA-*LEU*-ALA-*VAL*-VAL-*VAL*-TRP-*LEU*-X_2_-*LEU*-TRP-*LEU*-TRP-ethanolamine (with l-amino acids given in standard font and d-amino acids given in *italics*) (4). In gramicidin A, B, and C respectively, the X_1_ is VAL, VAL or ILE, and X_2_ is TRP, PHE, or TYR. The alternating l- and damino acids are crucial for the peptide to assume a *β*-helical form. This l- and dalternating motif is unusual for naturally occurring peptides (5); gramicidin is synthesised by non-ribosomal multi enzyme complexes, and the conversion of l-to damino acids takes place during the biosynthesis. At 16 residues, the sequence is too short to span the full hydrophobic lipid bilayer, however upon assumption of a helical dimer within the bilayer, conductance of monovalent cations occurs within the dimerised channel (6). Conductance occurs only when the N-termini of both of the dimers are linked via a hydrogen bond and stabilised within the membrane environment (7). The involvement of the N-termini in dimerisation was investigated through comparing the effect of C-terminal and N-terminal charged analogues (5). N-terminal charged analogues did not form conducting channels within black lipid membranes, which are model membranes developed in the 1960s, which are favoured for their ability to reconstitute proteins into the bilayer (8). Conversely, C-terminal charged analogues did produce conducting channels in these black lipid membrane environments. Gramicidin’s selectivity for monovalent cations (9) lead to gramicidin being a popular choice to study ion channels within lipid bilayers.

The helical dimer forms a cylindrical tube with twelve parallel *β*-sheet-like hydrogen bonds between residues in adjacent turns, which provide stability to each of the monomers (10). The hydrophobicity and structure of amino acid residues contribute further to the stability, with TRP at the membrane-solvent interface, and ALA and VAL buried more deeply. The abundance of TRP at the lipid-water interface and its abundance in membrane proteins suggests it plays an important anchoring role, with its favoured location driven by hydrogen-bonding between the indole N-H hydrogen and the carbonyl and phosphate oxygens of the local lipid headgroups (11). The gramicidin pore is ≈ 4 − 5 Å in diameter with a length of 26 Å and 6.4 amino acid residues per turn (12) (13), and typically does not have side on contact between adjacent helices (13). Outside of a lipid membrane environment, a pair of gramicidin peptides energetically favour a double stranded helix form (14) -such a structure is non conducting in a bilayer.

The *β*-helix structure is diverse in nature -they can contain several *β*-strand regions linked by turns (15), or follow the helical dimer motif that gramicidin is known for. Gramicidin’s propensity to form an ion channel for monovalent cations means its structure and sequence are powerful tools for the design of antimicrobial agents in the fight against multi-drug resistant bacteria. Gramicidin A’s potency as a disruptive ion channel to begin the process of apoptosis has seen its possibility as an anticancer agent explored (16).

A selection of peptides are discussed here, with the effects of different amino acid compositions and sequence lengths on the stability and flux of these peptides considered. Gramicidin dimer channels, in a concentration of 0.1 M NaCl with 100 mV applied across the membrane, have a conductance of a is 5.8·10^−12^ Ω^−1^ at 20°C (7). This is a benchmark for which the systems explored in this investigation will compared to.

An applied electric field was applied to measure the conductance of ions, as opposed to their passive diffusion across the membrane. This methodology comes with some complications and extra care must be taken with such a simulation. Research has found that the effect of electric field on interactions between ions and water is small (to the order of a few percent) for electric fields below 500 mV nm^−1^ (**?**). In research conducted using older force fields and parameters, it was notable that electric fields slightly reduce the conductance of Na^+^ and slightly increase the conductance of Cl^−^ (**?**). Electric fields of the order that was studied here have been shown to perturb the protein structure through localized dipole alignments (**?**), which in this case would be a beneficial artefact as it is desirable to have the helix aligned tightly to the z-axis. It has also been observed that electric fields with a strength of the order of that studied here has no effect on the *R*_*G*_ and RMSD of protein structures (**?**), suggesting that the largest challenge with this methodology will be in any artefacts introduced by the methodology with respect to the PME. These artefacts are more significant with longer simulations and larger electric field strengths, so this was addressed by maintaining an electric field strength that is low enough to have a minimal effect. A study on the effect of constant electric field methods in MD concluded that non-equilibrium properties are sensitive to the finite size of the simulation box, as the resistance and long-range dissipative effects are modified by the electric field potential (**?**), and recommends that systems are of sufficient size to combat this.

There are several design criteria that must be taken into account when designing a gramicidin-like peptide: (a) it must take on the form of a *β*-helix, which requires alternating land damino acids along its sequence, with h-bond interactions between the C=O and N-H groups driving the stability and structure of the peptide; (b) the outside of the core region must be highly hydrophobic, possibly with stabilising tryptophans at either end of each sequence, to stabilise the structure within the membrane; (c) it must permit the passage of water and ions through the membrane, enforcing a minimum pore width. As with any design, it is pertinent to only introduce complexity once the fundamentals are understood. Thus the sequences studied here are pure *β*-helical structures and will not contain loops, turns or *α*-helical sections as can often be found in *β*-helical proteins (17) (18) (19).

## 2 MATERIALS AND METHODS

### 2.1 Molecular dynamics simulation procedure

Unbiased molecular dynamics simulations were performed using the CHARMM36m forcefield (20) and GROMACS 2020.4 (21) (22). The membrane was composed of POPG and PVCL2 lipids in a 50:50 weight ratio (corresponding to a 1.8:1 numerical ratio) to mimic the membrane of *streptococcus pneumoniae* (23). Electrostatic interactions were computed using the particle mesh Ewald method (24), and a cutoff of 16 Å was used for the van der Waals interactions. The system was equilibrated in four stages with different time steps (0.1 fs, 0.5 fs, 1 fs, 2 fs), with the first two equilibration stages utilising an NVT ensemble, and the final two equilibration stages utilising an NPT ensemble. The production runs were performed in an NPT ensemble with a 2 fs time step. Each system had a total equilibration time of 50 ns and a total production run of 1 *µ*s. Frames were recorded every 0.1 ns. During the equilibration, a Berendsen thermostat (25) at 310 K was employed. Neighbour lists were updated every ten steps. In the production runs, the solvent, lipids, and protein were separately coupled to a heat bath at a temperuate of 310 K with a time constant *τ* = 0.1 ps using the Nosé-Hoover thermostat (26). Atmospheric pressure of 1 bar was maintained using the Parrinello-Rahman barostat (27) with semiisotropic (xy and z) pressure coupling with compressibility *κ*_*z*_ = *κ*_*xy*_ = 5 · 10^−5^ bar^−1^ and a time constant *τ* = 1 ps. The system was subject to a constant external electric field with *E*_0_ = +100 mV nm^−1^, applied in the z-direction. Analysis was performed using Python3.7, VMD (28), MDAnalysis (29), LiPyphilic (30), and PyMOL (31).

Ionic current was measured by tracking the movement of ions through the pore. This was done by utilising a custom script in Python. Ions were only counted as having passed through if they could either be counted inside the pore, or in consecutive frames were counted have been at the entrance on either side. Exceptions were added to the script to not count an ion as crossing if it approached the pore, sat there and then passed through the PBC to the other side. The same ion was not counted as crossing again if this occurred within 50 ns. Conductance was calculated using

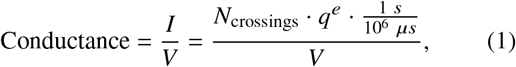

where *N*_crossings_ is the number of ions that cross the membrane in the full 1 *µ*s simulation, *I* is the current which was calculated by considering the number of crossings per second, multiplied by the unit charge, *q*^*e*^, and divided by the potential difference, *V* across the membrane. The 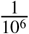 factor is for unit correction. This is an accurate measure of current provided the calculation of *N*_crossings_ is done carefully.

The angles that the centre of mass of the full sequence and the centre of mass of the upper and lower loops (defined as the first and last 8 residues respectively) made with each other, relative to if centre of mass of these upper and lower loops were positioned perfectly along the z-axis, were used to define the helical tilt. This served as a useful measure to determine how the inner hydrophobic residues of the helical sequence were being exposed to the lipid-water interface, as at larger angles it was more deviated from the z-axis, which positions the hydrophobic (ALA, LEU, or VAL) residues closer to the lipid-water interface than they would naturally prefer to be. This is show diagrammatically in figure 1.

**Figure 1:**
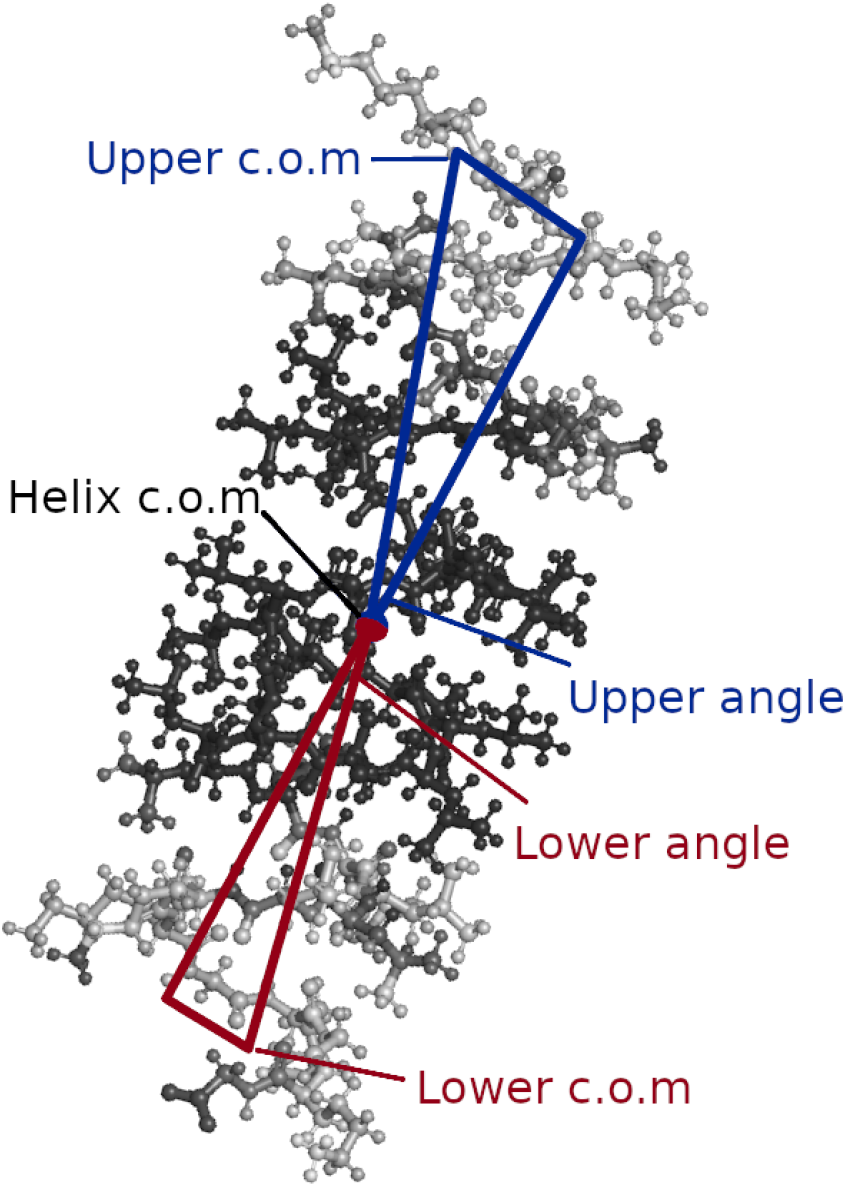
A diagram to demonstrate the trigonometric set up to calculate the angles made between the c.o.m of the helix and it’s upper and lower extrema. These are the angles plotted in figure 6.

### 2.2 Molecular dynamics preparation

The *β*-helix design began from measuring the *ψ* and *ϕ* angles of an experimentally resolved structure of gramicidin A (pdb: 1GRM) (32), using VMD. These values were averaged and used as a template for the initial structures. Using hippo/Atomix^1^ it is possible to generate amino acids minimised with specified dihedral angles. Using the exact averaged value from 1GRM yielded an unstable structure that at residue counts of <60 amino acids did not fully span the membrane environment. This structure was used as a basis, and then the dihedral angles were modified by ±1°. This was repeated 40 times, with each result saved to a multi-frame .pdb file (figure 2). The overall sequence and length could then be analysed and visually examined. As the dihedrals were increased, there was only a marginal change in pore size, but the C=O:N-H distance increased, as well as the end-to-end peptide length.

**Figure 2:**
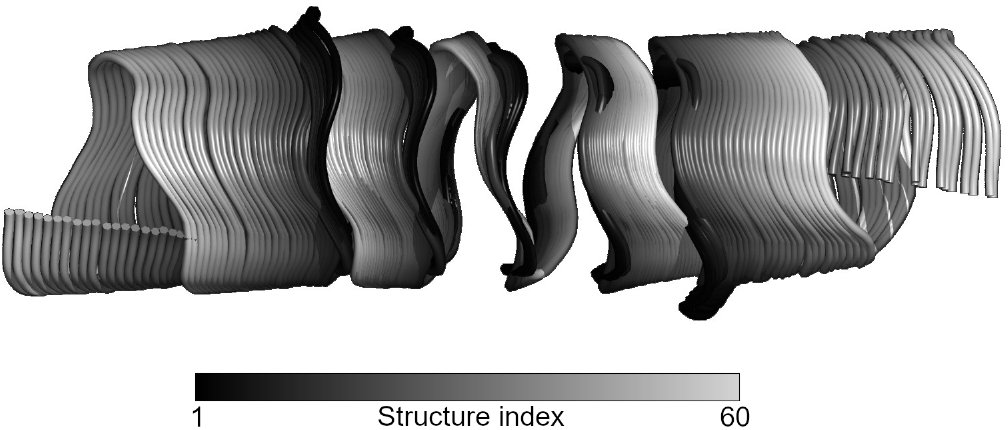
The unfolding *β*-helix shown by its backbone with each frame a separate colour, with the starting stracture in black and the lightest grey the final structure. The gradual increase in length and C=O:N-H bonding is evident.

The original design used GLY residues to span the hydrophobic membrane, but it became apparent that an amino acid with greater hydrophobicity was required. A GLY helix with LYS and d-TRP at the ends proved to unravel and “unclick” - the crucial C=O:N-H bonds shifted and stretched, the “unclicking” leading to the new positioning of these constituents atoms out of step with each other. Backbone interactions along the coil of the helix were not sufficient to maintain its shape. Notably, a pure GLY helix without LYS and dTRP at the ends more unstable. The result was two unaligned *β*-helices, permitting the entry of solvent into part of the bilayer, and a unstructured strand of GLY connecting to the two helices. Whilst this may prove to be sufficient to destabilise the membrane, this did not permit a continuous flow of water and ions through the membrane environment. Instead, it opened up space for a pocket of water to exist within the membrane, further unravelling the helix, but with no conductance.

The original GLY designs also proved useful for providing estimates for a range of sequence lengths to cover in the next stage of the design - sequence lengths of less than 40 proved insufficient as one end of the helix was liable to obfuscation by the lipid headgroups. Conversely, sequence lengths above 60 became overly tilted and were more liable to “unclicking”, as well requiring wider simulation boxes to account for their tilt, increasing calculation times. These original pure GLY helices served an effective initial design stage. It provided a minimum and maximum sequence length to test, as well justify the choice of LYS and d-TRP to be the membrane-solvent interface residues.

Previous studies have elucidated the propensity of amino acid sequences (33) to sit at certain heights within the membrane. These results were used to inform the post-GLY *β*-helix design. ALA, LEU, and VAL were all evident as being the best possible candidates for these structures. They energetically favour the centre of the hydrophobic bilayer, and are less likely to found closer to the interface (33). This propensity of their character was expected to lend to a more stable structure within the membrane environment. It also provided a range of different side chains which were hypothesised to offer some stability within the membrane as they would have intearctions with the lipid tails. The presence of these residues within the gramicidin sequence supports this as a sensible approach.

To permit more favourable like-for-like comparisons, each *β*-helix was composed of only one species of amino acid (with both l- and dchiralities) in its core, and with the first eight and last eight residues of each helix containing alternating LYS and GLY, or LYS and d-TRP. It was expected that a charged amino acid like LYS would enhance headgroup interaction and provide a stablising effect to the structure - as was observed with the pure GLY helix originally tested. Indeed, previous studies have found for gramicidin mutants that using amino acids with cationic side chains in their sequence (specifically LYS) can reduce hemolytic activity whilst retaining antimicrobial potency (34). The four d-TRP residues at either end were chosen due to their presence in the gramicidin sequence and their known capacity to lie at the membrane-solvent interface due to their side chain indole (35) (36).

Figure 3 shows the conductance of the 66 systems examined in this study. Ionic conductance was considered only if the ion did not recede from the same side of the pore for 10 consecutive ns; the ion was also required to be measured at both sides of the pore during this window. In the instance that this criteria was not met, the ion was not considered in the calculation for conductance. Twelve of the 66 systems showed a conductance ≥ 30% of the gramicidin conductance. This value was chosen as as it yields a conductance value to the same order of magnitude as gramicidin, and it served a neat way to delineate between systems which performed averagely and those which performed well. For example, decreasing the threshold to ≥ 10% of the gramicidin conductance doubles the number of candidate systems to 24. This doubles amount of data to explore, but the additional data this lower threshold brings is of systems that performed more poorly. Thus, these twelve systems were then considered the candidate systems and examined in further detail - the full list of candidates in available in table 1. For these candidates - three of the twelve have d-TRP residues at the ends of their sequences, four of the twelve have VAL for their core residues, six of the twelve have ALA for their core residues, and two of the four have LEU for their core residues. Thus, across all the candidates, all the different components that made up the helix could be evaluated.

**Table 1:** Candidate systems explored in detail, with their name and full sequence. l-amino acids are shown in upper case and D-damino acids are given in lower case.

| | Candidate name | Candidate Sequence | Conductance ($\Omega^{-1}$) |
| --- | --- | --- | --- |
| 1 | 40GVG | KGKGKGKG-[Vv]<sub>12</sub>-KGKGKGKG | $(2.51 \pm 1.62) \cdot 10^{-12}$ |
| 2 | 42WAW | KwKwKwKw-[Aa]<sub>13</sub>-KwKwKwKw | $(1.79 \pm 1.76) \cdot 10^{-12}$ |
| 3 | 44GAG | KGKGKGKG-[Aa]<sub>14</sub>-KGKGKGKG | $(2.48 \pm 1.53) \cdot 10^{-12}$ |
| 4 | 44GVG | KGKGKGKG-[Vv]<sub>14</sub>-KGKGKGKG | $(2.09 \pm 1.52) \cdot 10^{-12}$ |
| 5 | 44WLW | KwKwKwKw-[Ll]<sub>14</sub>-KwKwKwKw | $(1.79 \pm 1.55) \cdot 10^{-12}$ |
| 6 | 46GAG | KGKGKGKG-[Aa]<sub>15</sub>-KGKGKGKG | $(3.56 \pm 1.69) \cdot 10^{-12}$ |
| 7 | 46GLG | KGKGKGKG-[Ll]<sub>15</sub>-KGKGKGKG | $(2.02 \pm 1.83) \cdot 10^{-12}$ |
| 8 | 48GAG | KGKGKGKG-[Aa]<sub>16</sub>-KGKGKGKG | $(2.58 \pm 1.84) \cdot 10^{-12}$ |
| 9 | 48GVG | KGKGKGKG-[Vv]<sub>16</sub>-KGKGKGKG | $(1.68 \pm 1.96) \cdot 10^{-12}$ |
| 10 | 50GAG | KGKGKGKG-[Aa]<sub>17</sub>-KGKGKGKG | $(2.12 \pm 1.80) \cdot 10^{-12}$ |
| 11 | 58GVG | KGKGKGKG-[Vv]<sub>21</sub>-KGKGKGKG | $(2.14 \pm 1.87) \cdot 10^{-12}$ |
| 12 | 60WAW | KwKwKwKw-[Aa]<sub>22</sub>-KwKwKwKw | $(1.92 \pm 1.66) \cdot 10^{-12}$ |

**Figure 3:**
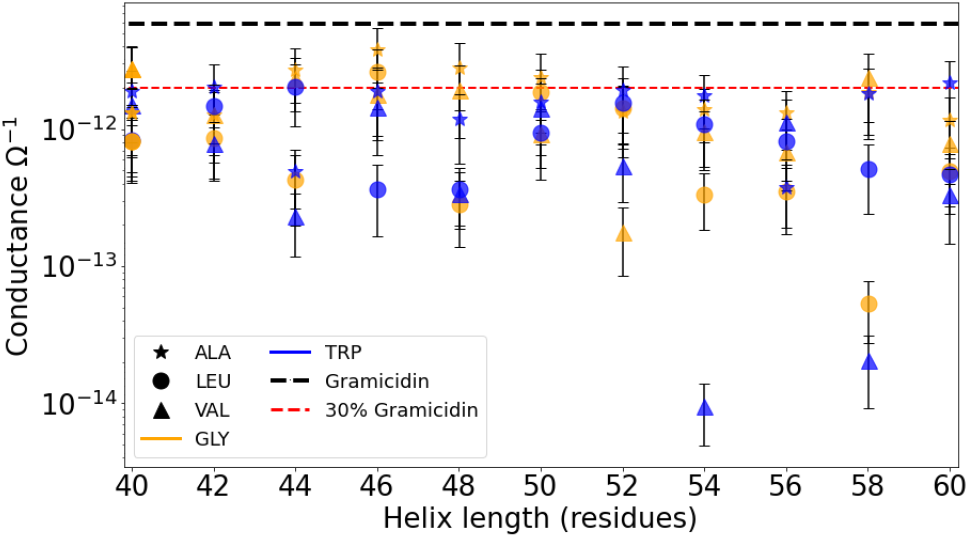
Conductance values for each system. The black dashed line shows the value of gramicidin’s conductance as per literature, with the red dashed line showing 30% of this value. Any point on or above the red dashed line is of the same order of magnitude of gramicidin and considered a good candidate. The error bars on this plot were calculated by dividing each simulation up into 100 ns windows, which yields 10 conductance values per system. The standard deviation of these 10 values is what is shown here. Systems with lower overall conductance had larger standard deviations due to having a higher proportion of low/no conductance samples, so this method is a little crude but effective for our purposes.

## 3 RESULTS

There is a degree of unravelling that occurs at the membranesolvent interface in the outer residues of the helix. This was observed in all systems, although it was more pronounced in longer sequence lengths and in systems without TRP present at the water-lipid interface. What this unravelling looks like in practice can be seen in figure 4, where one terminus of the protein sequence has lost its helical shape and presents as unstructured loop. This has the effect widening the pore at this end of the helix, increasing access to the pore for water and ions.

**Figure 4:**
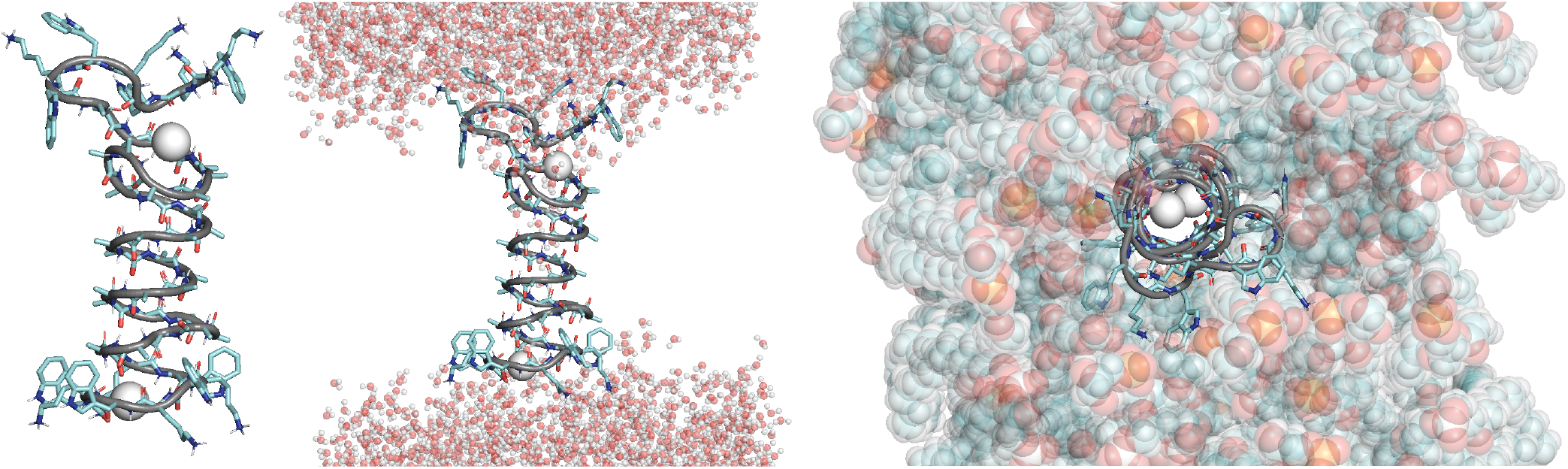
The 48-ALA-TRP system in three different views. Carbons are shown in cyan, nitrogens in blue, oxygens in red, and polar hydrogens in white, all as a ball and stick representation, but with the helical backbone highlighted over the top as a grey cartoon ribbon. The K+ ions are shown as white beads. **Left:** Side on, with membrane and waters hidden and two K+ ions shown in the pore. **Middle:** Side on, with waters shown. **Right:** Top down, with waters hidden and the membrane shown, with phosphorous in orange and other atoms coloured as they are for the helix. Despite the unravelling of upper loop, the pore and conductance remains stable.

RMSD (root-mean square deviation) is a quantitative measurement that evaluates how similar two structures are by comparing their atomic positions. Thus, RMSD distributions are a good measure of how much the helix unravelled - narrower and smaller values for the peak of the distribution indicated that a helix was more prone to keeping its shape.

The RMSD distributions in figure 5 are shown with solid lines for helices containing tryptophan and dashed lines for helices containing glycine. 42WAW, 44WLW and 46GAG shown the leftmost distributions and the narrowest distributions. The presence of both of the shorter conducting tryptophan helices is not a coincidence, and is a direct consequence of the role tryptophan plays in stabilising the helix through its propensity to sit at the membrane-solvent interface. Unravelling is exacerbated by the pore as water is able to enter pockets of space opened by the flexibility of the helical loop, and with the water in place these atoms are displaced and no longer available to take part in structure-stabilising backbone interactions. As interactions with the lipid headgroups decrease the helical backbone H-bond interactions, the strand loses its *β*-helical structure and more closely resembles a random-coil; stochastically this can lead to the charged LYS side chains blocking the entrance the pore. The residues that are enveloped by the hydrophobic environment of the lipid tails remain tightly coiled and retain their helical shape.

**Figure 5:**
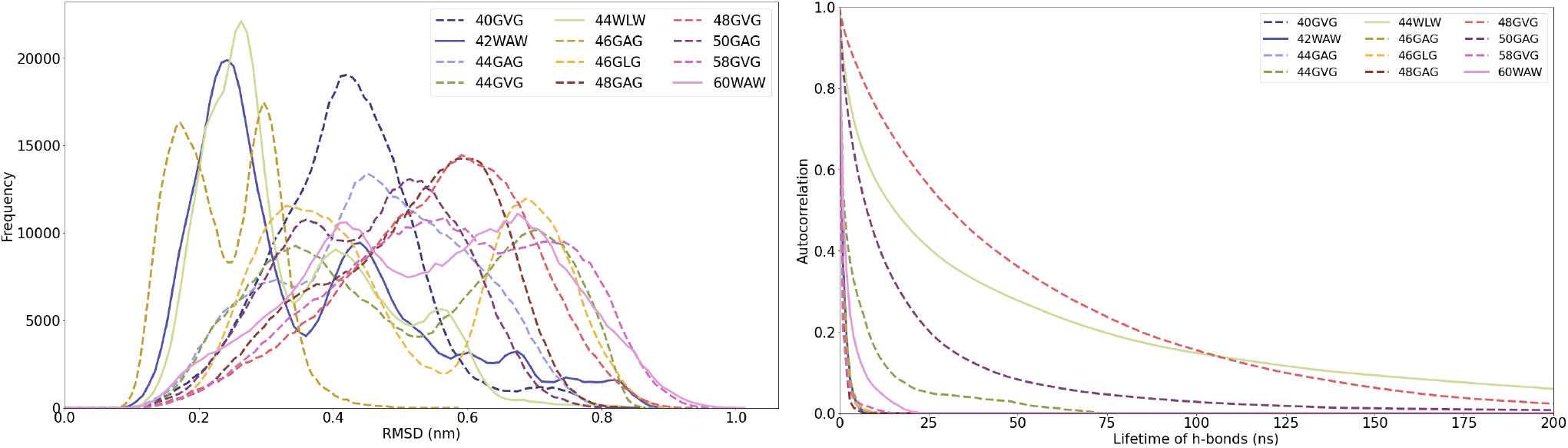
**Left:** Root mean square deivation (RMSD) profiles for the twelve conducting systems. The further to the left and narrower a peak is, the better the *β*-helix holds its shape. **Right:** Autocorrelation of the hydrogen-bond lifetimes for the twelve conducting systems. A *β*-helix’s backbone retaining a tighter coil, and more/better C=O:N-H coupling along the backbone, can be identified by a greater area underneath the curve of the autocorrelation function.

The autocorrelation function of H-bond lifetimes (righthand side of figure 5) reveal which sequences maintain C=O:N-H bonds along the rise of the helix. 48GVG, 44WLW, and 50GAG are the best performers by this metric. Given that 44WLW possesses a narrow RMSD it is expected that this is a relatively fixed structure, despite existing in a highly interactive environment, particularly for side chains that have favourable interactions with chemical groups in the lipids. 20% of its hydrogen-bond interaction pairs spend at least 90 ns within 3.3 Å, per interaction. Many of these backbone interactions are constantly forming and reforming and thus will exist for longer than this lifetime of 90 ns, with only brief interruptions in this contact. How the helix sits within the membrane can be evaluated by considering the angles made by the ends of the helix with the centre of mass (c.o.m) of the helix itself, and the profile of interactions between helix atoms and the lipid headgroups. These profiles are captured by using the contact analysis function of MDAnalysis, and colouring each atom according to the frequency at which it contacts the lipid headgroups relative to other atoms in the helix (37). These figures are shown in figure 6. They are concise but a lot can be learned about these sequences from this coloured contact analysis. The upper and lower angles, also shown in figure 6 and defined in figure 1, are typically coupled for each system, following the same trajectory despite differences in their range of values. Generally the upper leaflet angle is smaller, but this value is influenced by the unravelling typically occurring on the upper leaflet strands. Why the helical unravelling occurs only at the upper leaflet is unclear, as there is no biasing of the systems to do so and the helix is centred with the bilayer, so it should extend an equal distance above and below the centre. The value for the angle by either leaflet rarely exceeds 20°. This tilt angle increasing means that end of the helix is tilting further away from the membrane normal, which in turn influences what parts of the hydrophobic core of the helix are exposed to the headgroups, with a large value exposing the hydrophobic side chains of the core to the lipid headgroups.

**Figure 6:**
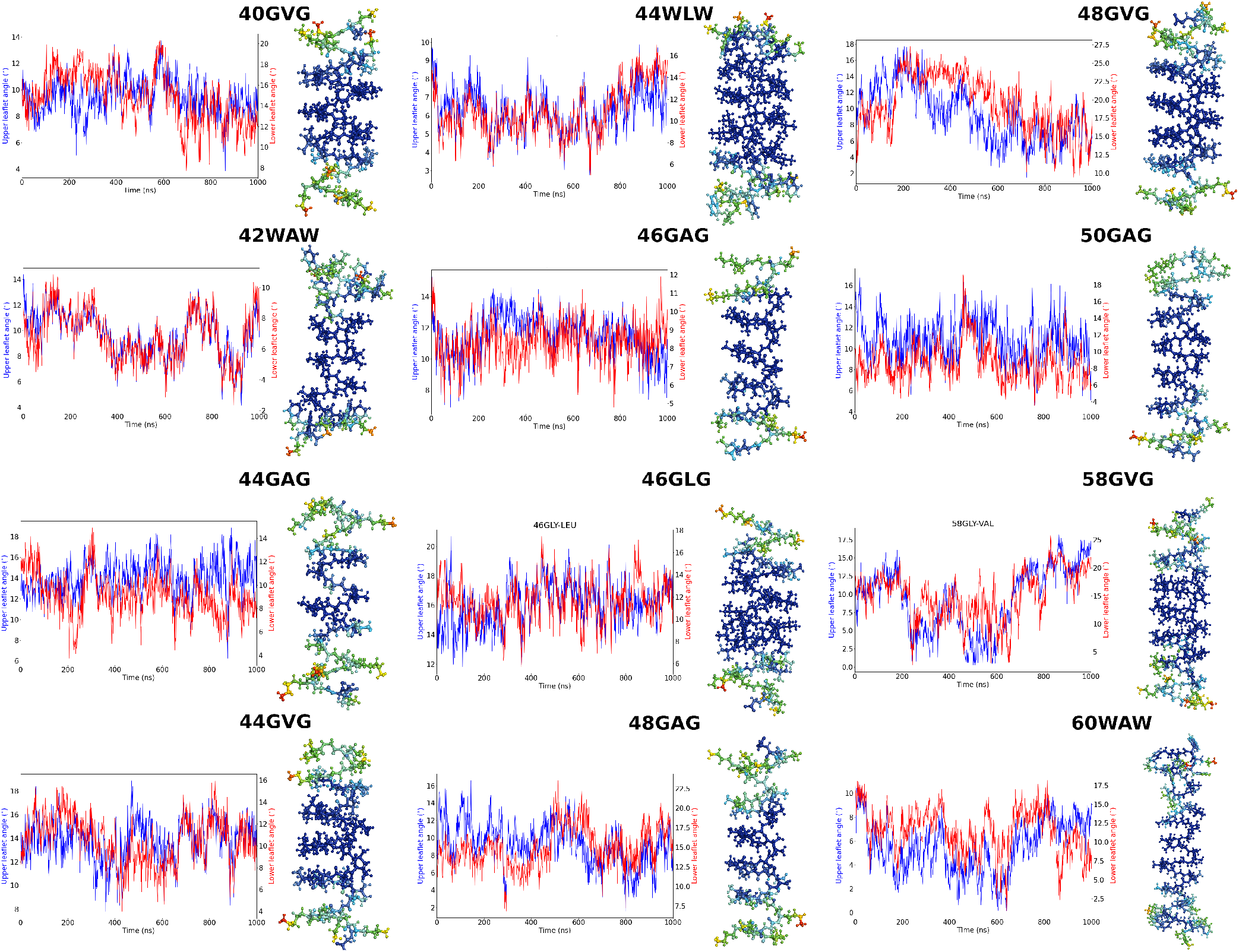
The angle between centre of mass (c.o.m) of the helix and the end loops at the upper leaflet (blue) and lower leaflet (red) are shown, along with headgroup contact analysis for all-atoms in the helix. Headgroups were considered to be anything above the glycerol unit (with tails considered to be below the glycerol unit). Atoms in blue have zero contact with these components of the lipid, and red atoms have the maximum contacts for that system. A contact was defined as being within 3.3Å. The helix atoms are coloured according to the scale indicated, with the maximum contact for each system in a red hue, and the blue atoms having zero contacts.

It is notable that for longer sequences, it is common to find interactions with the headgroups a full helical turn toward the center of the helix. In each system interactions with the headgroups are driven by LYS side chains, but the longer sequence length causes ample exposure of the beginning of the hydrophobic core of sequence to the lipid-water interface. Such an effect is less pronounced in LEU helices, a result which can be attributed to the interactions between the LEU side chains and the lipid tails. All else being equal, the hydrophobic core of VAL helices are less exposed than their ALA counterpart to the lipid headgroups, particularly at shorter sequence values. This is likely driven by the interactions between the VAL side chains and the lipid tails. With the tryptophan residues, half of the indole experiences headgroup interactions, whilst the other half does not. This is in agreement with what we expect from tryptophan’s orientation at membrane-solvent interfaces (35) (36).

The interaction of these deeper residues with the membrane headgroups can be partly understood by considering the RMSD, shown in figure 7. It is generally true that the magenta line, which shows the average RMSD of the first and last four residues over the full simulation, is both the noisiest curve and possesses the largest value. The difference between the RMSD of the magenta line and the black line (which shows the middle residues) is generally less pronounced for helices containing d-TRP at its ends. This result is likely driven by TRP’s propensity to sit at the membrane solvent interface, as in the other helices it has been replaced by GLY which has no such favourable interactions to help anchor it. Large, instantaneous changes in the RMSD often (but not always) corresponds to the passage of an ion, as the the close approach of a charge has a strong electrostatic interaction with LYS present in all of the helices. RMSD serves as an efficient proxy for a measure of the unravelling and the helices with their Nand Cterminal residues which are more prone to ‘spaghettification’ show a larger RMSD.

**Figure 7:**
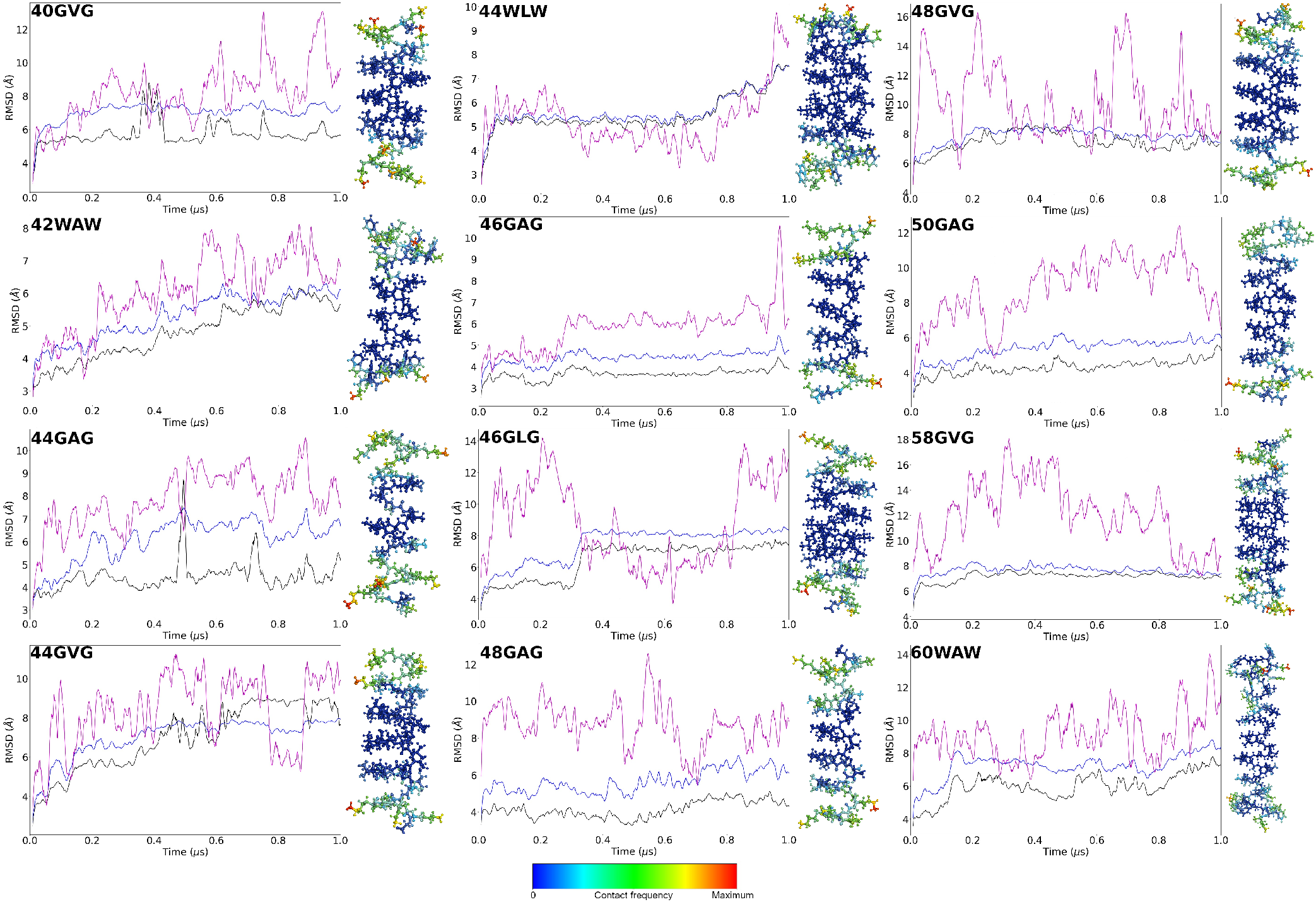
The RMSD profile for all atoms of the helix (blue line), the first 4 and last 4 residues of the helix (magenta line), and the ‘centre’ of the helix (black line). We define the center of the helix to be the residues numbered 4 to *n*-4 (all but the first and last 4 residues). Accompanying each RMSD plot to its right is headgroup contact analysis for all-atoms in the helix. Headgroups were considered to be anything above the glycerol unit in the lipids (with tails considered to be below the glycerol unit). Atoms in blue have zero contact with these components of the lipid, and red atoms have the maximum contacts for that system. A contact was defined as being within 3.3Å. The helix atoms are coloured according to the scale indicated, with the maximum contact for each system in a red hue, and the blue atoms having zero contacts.

The width of the pore is critical for the peptide’s function. The HOLE program (38) (39) (40) was utilised to measure the width of the pore across the membrane at each point along the z-axis, as well as provide a measure of the standard deviation of such a value, in figure 8. An optimal helix will have a very small standard deviation at every point within the membrane, demarcated by the black lines. The two shorter tryptophan distributions (42WAW and 44WLW) show at certain points neglible *σ* values which reveals a remarkably stable structure at that section of the membrane, with little to no movement. This behaviour is lost for the longer 60WAW tryptophan helix, suggesting there exists an optimal sequence length between 42 and 58 amino acids. It is generally true for the GLY capped helices too, that longer sequences showed a greater deviation in their width. For the shorter helices, deviations within the bounds of the membrane are somewhat explained by the loss of structure of the end helical loop and the breaking of hydrogen-bond interactions as a result of this, leading to a wider structure with less positional rigidity. Regardless of sequence length, there is an approximate uniformity of 2.8-3.5 Å for the pore radius, for the sequence that resides inside the membrane environment fully. The pore radius increasing to values ≥ 4 Å before the line denoting the membrane interface is a sure sign that the system has unravelled within the membrane environment. Considering the aim of this research project, a stable pore structure is essential, and so this remains a key benchmark for determining the efficacy of the structures. As such, a sequence length of 48 looks to be a effective minimum length to pursue in future iterations of this work, given it effectively meets this criterion.

**Figure 8:**
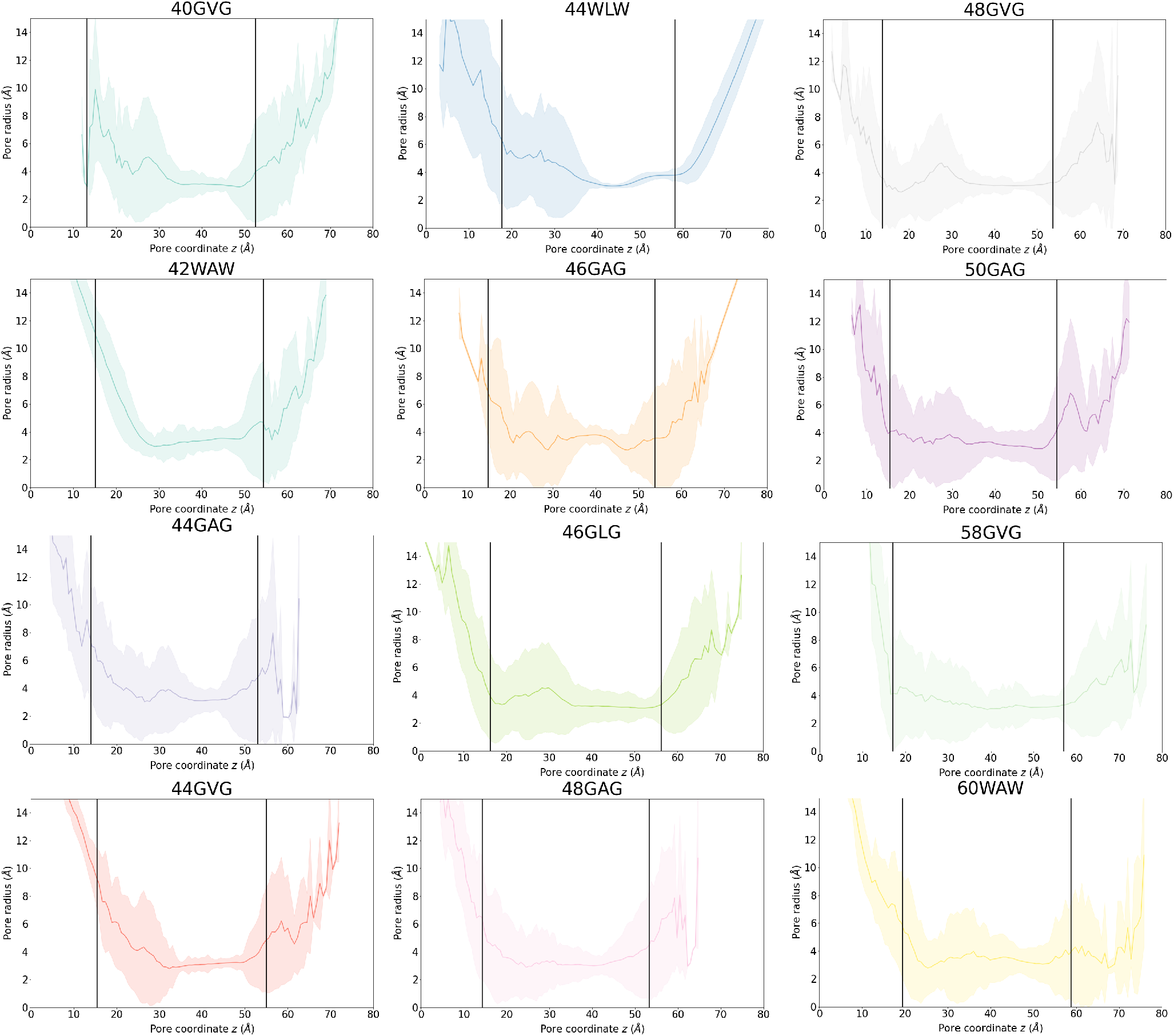
The pore width for each of the conducting helices, calculated using the HOLE module within MDAnalysis. The solid coloured line shows the mean pore width (*µ*_HOLE_) over the simulation for a given coordinate along *z*, and the shaded area shows the standard deviation, *σ*_HOLE_, of this mean. The black solid lines indicate the bounds of the membrane bilayer.

## 4 CONCLUSION

These *β*-helical peptides show promise as candidate peptides for antimicrobial peptides in Gram-positive lipid membranes. There are several avenues to pursue for improvements to such peptides; one such approach would be to use nonhomogeneous protein sequences. Given that ALA made up six of the 12 candidates, but showed the greatest deviation in pore width, combining ALA and VAL at the appropriate points along the amino acid sequence may result in a helix that offers a consistent width and can efficiently conduct ions. VAL would be a fine choice due to the lower standard deviations seen in the 44GVG and 48GVG systems within the membrane environment. While a small deviation is also observed in the 44WLW systems, LEU’s relative scarcity suggests it is less effective. More residues per turn would afford a wider pore, although a higher sequence length would make synthesis for future in vivo testing more challenging. However this wider pore would also allow more than just water and ions to passage through the pore, and could potentially be a way to allow select small molecules into the cell, which would be a fine result. Another direction would be attached lipids at the N and C termini of the helix to offer stability. This last method may also result in greater binding affinity to membranes, and can help fine tune the selectivity of the helix for particular membrane compositions through the choice of attached lipid. Unfortunately, none of the tested sequences proved to be as potent as gramicidin, but this project serves as an important first step in the right direction. This remains an open research project with the original aims not fully achieved, but this is an important step towards solving this problem. With the process of selecting the sequence, preparing, and running the MD simulations being automated, this work is in a good place to be further developed.

## 5 AUTHOR CONTRIBUTIONS

MBU designed the research. EL generated the structures, performed the analysis, and wrote the article. MIW guided and supported the research. MBU reviewed the article.

## 6 ACKNOWLEDGMENTS

This work was done with funding from EPSRC via the Cross-disciplinary Approaches to Non-Equilibrium Systems (CANES) CDT and the BBSRC. Simulations were carried out on the GRAVITY and CREATE supercomputing infrastructure at King’s College London. The authors have no relevant financial ties to disclose. The authors thank Dr. Dominique de Jong-Hoogland for her sage advice and her help in reviewing the article.

## SUPPLEMENTARY MATERIAL

Made available on the lead author’s github.

## Footnotes

1 Developed and maintained by Martin and Jakob Ulmschneider, available at https://www.biowerkzeug.com/

## Notes

### Competing Interest Statement

The authors have declared no competing interest.

### Summary of Updates

Added an additional author, updates to text to improve readability.

